# Notions of similarity for computational biology models

**DOI:** 10.1101/044818

**Authors:** Ron Henkel, Robert Hoehndorf, Tim Kacprowski, Christian Knüpfer, Wolfram Liebermeister, Dagmar Waltemath

**Affiliations:** Heidelberg Institute for Theoretical Studies, Heidelberg, Germany; Computational Bioscience Research Center Computer, Electrical and Mathematical Sciences & Engineering Division King Abdullah University of Science and Technology Thuwal, Kingdom of Saudi Arabia; Department of Functional Genomics Interfaculty Institute for Genetics and Functional Genomics, University Medicine and Ernst-Moritz-Arndt University Greifswald Greifswald, Germany; Institute for Computer Science, University Jena, Jena, Germany; Institute of Biochemistry, Charité – Universitätsmedizin Berlin, Berlin, Germany; Department of Systems Biology and Bioinformatics, Institute of Computer Science, University of Rostock, Rostock, Germany

**Keywords:** Systems biology, Network model, Model versioning, Information retriaval, Systems Biology Markup Language

## Abstract

Computational models used in biology are rapidly increasing in complexity, size, and numbers. To build such large models, researchers need to rely on software tools for model retrieval, model combination, and version control. These tools need to be able to quantify the differences and similarities between computational models. However, depending on the specific application, the notion of “similarity” may greatly vary. A general notion of model similarity, applicable to various types of models, is still missing. Here, we introduce a general notion of quantitative model similarities, survey the use of existing model comparison methods in model building and management, and discuss potential applications of model comparison. To frame model comparison as a general problem, we describe a theoretical approach to defining and computing similarities based on different model aspects. Potentially relevant aspects of a model comprise its references to biological entities, network structure, mathematical equations and parameters, and dynamic behaviour. Future similarity measures could combine these model aspects in flexible, problem-specific ways in order to mimic users’ intuition about model similarity, and to support complex model searches in databases.

## 1 Introduction

“Over the past few decades, mathematical models of molecular and gene networks have become an important part of the research toolkit for the biosciences” [1]. Mathematical models are formal representations of natural systems that can help answer questions about the complex system they represent [2]. According to Robert Rosen, a model establishes a *modelling relation* between a formal and a natural system: the formal system encodes the natural system, and inferences made in the formal system can be interpreted (*decoded*) as statements about the natural system [3]. Computational models in biology serve as abstractions of biological systems. Biochemical models, for example, associate model components, such as mathematical expressions, objects, or variables, with biochemical entities such as molecule species or chemical reactions. Depending on the scientific questions addressed and on the available data, a biological system may be described by models of different scopes and levels of granularity, reflecting different views of the system.

Computational models can be based on a number of mathematical formalisms [1]. Here, again, the choice of a particular approach largely depends upon the type of question asked and on the data available [2]. Metabolic and signaling pathways are usually modelled by ordinary differential equation systems (ODEs), and the resulting models are known as kinetic models. Larger metabolic systems are typically described by constraint-based network models that capture stationary metabolic fluxes, but disregard enzyme kinetics. Gene expression dynamics can be modelled by kinetic models, stochastic processes, or discrete dynamic processes such as Boolean networks [4]. Spatial cell models may even involve partial differential equations (PDEs). In addition, the rise of synthetic biology and the development of virtual organisms require hybrid modeling approaches [1].

Models that are formally encoded in standard formats can be processed by and exchanged among different software tools. The Systems Biology Markup Language (SBML) [5] and CellML [6] are two XML-based de *facto* standards that encode the entities and interactions in biological models. Both enable software interoperability across the diverse landscape of modeling, visualisation, and simulation software [7]. To further standardise the representation of models and enable interoperability between them, the biological meaning of model components can be specified by semantic annotations that relate components (e.g., a variable representing glucose concentration) to common identifiers defined in ontologies or public databases (e.g., `CHEBI`:17234, which is the identifier for glucose in the ChEBI database [8]). Together, standard formats and semantic annotations foster the reuse of data in the biosciences [9]. In addition, the reuse of models across research groups and scientific use cases demand sophisticated model management strategies for storage, search, retrieval, version control and provenance [10].

For any domain, many notions of similarity can be used depending on the aspects between which similarity is determined. For example, to decide whether two persons are alike, we need to choose what features of the person to focus on, and we may assign priorities to selected features. Then, we may compare two persons by how tall they are, by the shapes of their faces, or by their behaviour – with very different results. Certain groups of people may be easily distinguished by size (e.g., children versus adults), while the same criterion will fail to predict other distinctions (left-handed versus right-handed people). Also in computer science, data objects can be compared with regard to different aspects depending on the purpose of the comparison. For example, when comparing image files, typical features are the colours used, the objects shown, or file size and type. The choice of the features and the choice of the similarity measure depends on the intended application.

The comparison of models is already implemented in a few software tools as part of their similarity search. For example, public model repositories such as the CellML model repository [11] can utilize a ranked retrieval system [12] that supports users in finding models relevant to their research. For such a ranked retrieval system, similarity measures are applied and adapted to yield a measure of model similarity based on various features of models [13]. Another example is semanticSBML [14], an online software tool that provides functionality for clustering, merging, and comparison of SBML models based on semantic annotations.

In this paper we overview and categorise existing similarity measures between models and their components. These measures may rely on aspects such as the biological entities described, network structure, model assumptions, mathematical statements and parameter values, or the dynamic behaviour displayed in simulations. We discuss the different aspects and show how they can be incorporated into computable similarity measures. We furthermore show applications for these measures, using the examples of model search, clustering and merging.

## 2 Formal notions of similarity for computational models

### 2.1 Similarity measures

The general notion of similarity can be conceptualised by mathematical functions called *similarity measures*. Intuitively, a similarity measure, for some type of objects, assigns to each pair of objects a similarity value. Larger values signify greater similarity. If properties of the objects are represented in some property space, similarity resembles an inverse distance: objects with small distances are considered similar, objects with large distances dissimilar. Similarity measures are often normalized to yield values between 0 and 1. A similarity σ = 1 then implies that two objects are identical (or indistinguishable) with regard to the properties considered, while entirely different objects will have a similarity *σ =* 0. Formally, a normalized similarity measure for a set *X* is defined as a function which assigns to pairs *x*_1_,*x*_2_ ∈ *X* a value *σ* ∈ [0,1]. This *σ* is called the “similarity value”. Most similarity measures are symmetrical and satisfy the triangle inequality.

### 2.2 Model similarity based on single aspects

Similarity measures for models refer to specific model aspects. For example, the similarity between two networks can be determined by aligning the networks and assessing their common overlap [15]. The result is a similarity value with respect to network structure. Alternatively, the similarity between two models can be calculated by comparing simulated time series [16], by comparing semantic annotations [12], or by identifying occurring patterns, etc. In the following, we define similarity *with respect to* certain model properties. We first define similarity with regard to a single aspect. Later in the paper, we also discuss similarity with regard to a combination of aspects.

To formally express similarity with regard to a single aspect, we introduce the projection of a model *M* onto an aspect *α*, for example, the projection of a dynamical model onto the underlying network topology. All other model features, which do not belong to this aspect, are ignored. The similarity measure between the projections of two models then determines the similarity of these models with respect to that aspect (see (Figure 1). More formally, for a model aspect *α* the α-similarity between two models *M*_1_*, M*_2_*, sim_α_(M*_1_*, M*_2_) is defined as σ_α_(Π_α_(*M*_1_), Π_α_(*M*_2_)), where Π_α_ is the projection of a model onto aspect α and σ*_α_* is a similarity measure for aspect α. This scheme applies both to models as mathematical objects and to encoded models, i.e., text files representing these models, and allows for statements such as “model 1 and model 2 are similar with respect to aspect α”. This definition may lead to undefined similarity measures, because we consider the projection function Π_α_ a partial function. If a model lacks the aspect for which the similarity measure is defined, the similarity to other models remains undefined as well.

**Figure 1.**
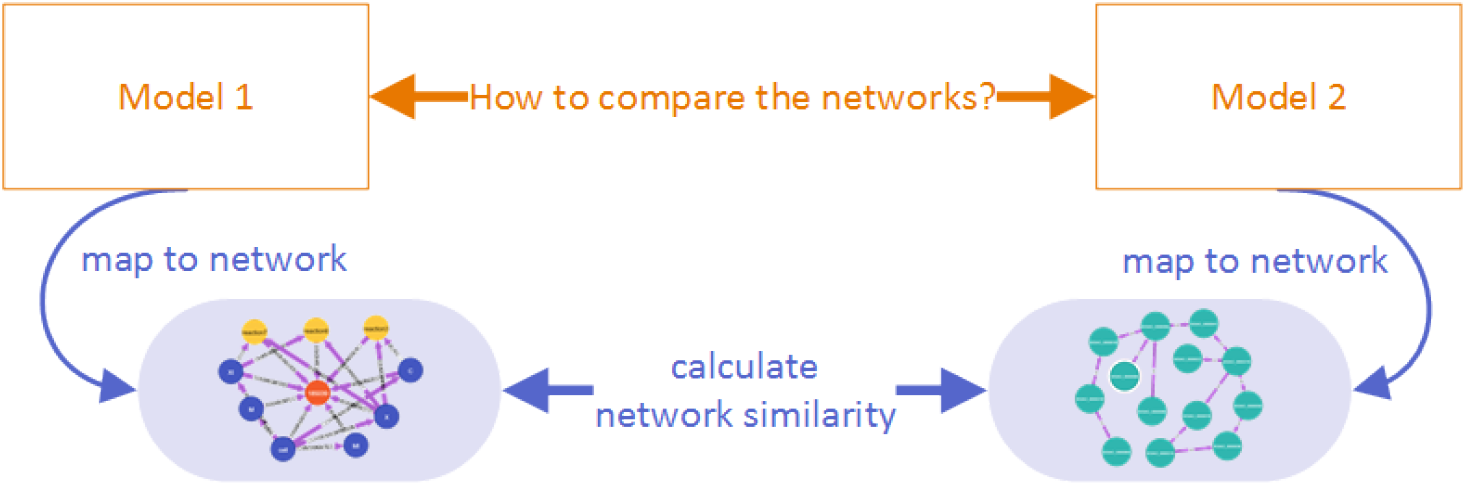
**Comparison of two models based on an aspect**. If two models are to be compared, the models’ properties are mapped to available aspects (their network properties in this case). The similarity of the aspect is calculated by measures appropriate for the aspect, e.g., for network similarity [15].

With the same scheme, models can also be compared to other types of data, e.g., to sets of experimentally measured compound concentrations. To see whether a model and a data set refer to similar sets of compounds, we could choose the sets of compounds as the aspect we’re focusing on. Mathematically, if a projection Π_α_*(M)* yields the aspect α of a model *M* and a related projection 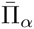 projects data sets *D* to the same aspect α, then an α-similarity function σ*_α_* can be used to find data sets that resemble a model *M* with respect to α, by computing 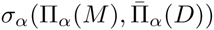.

### 2.3 Combined similarity measures

Similarities arising from different model aspects can be combined to define more complex measures. For example, two models may only be considered similar, if they describe similar biological entities *and* if they describe them by similar mathematical formulae. The combination of different similarity measures occurs frequently in model search, e.g., users looking for “models that describe aspects of the Cell Cycle and use similar parameter space”.

To perform such complex tasks, we need to aggregate multiple similarity scores into one single score. We can compose or decompose complex similarity measures by projecting models on individual aspects and then combining the resulting similarities between these aspects. Formally, we can express this as follows (without loss of generality limited to normalized similarity measures): 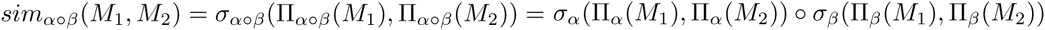 where o is a function o: [0, 1] x [0, 1] → [0, 1].

For larger numbers of properties to be combined, similarity measures can be conveniently defined by feature vectors. A feature vector represents different model properties by a list of numbers arranged in a vector. For example, the elements of the vector could be zeros or ones, denoting the occurrence of certain annotations in the model, or integer numbers denoting the frequencies of certain network motifs in the network. Once models have been translated into feature vectors, a variety of methods from multivariate analysis can be applied, including supervised and unsupervised classification [17]. Simple similarity measures for feature vectors can be defined based on Euclidean distances, the Jaccard index, or on normalised scalar products (i. e., the cosine of the angle between feature vectors). To weigh different features and to account for their known relationships, special metrics can be used, e.g., quadratic forms instead of simple scalar products.

## 3 Model comparison based on specific model aspects

Relevant aspects of models can be extracted (or calculated) from the encoded models. Depending on the goal of a comparison and on a model’s representation, the computable similarity measure is chosen. When models are presented as network graphs, it will be natural to compare them by network structure using a graph similarity measure. For example, the Systems Biology Graphical Notation (SBGN, [18]) for graphical display supports comparisons of network structural and visual similarity. If only model equations are given, network similarity would be more difficult to recognise even if all necessary information is implicitly given. For example, SBML and CellML for model structure and formulae support comparisons of structural and mathematical similarity. The SBO and other ontology-based annotation support semantic similarity comparisons. According to our framework, the projection function Π_α_ can be more or less complex to define (and evaluate) given a certain model representation. Consequently, certain similarity measures will be more natural given a certain model representation.

We propose to classify similarity measures based on five types of model aspects: (i) model encoding; (ii) biological meaning; (iii) network structure; (iv) mathematical statements and numerical values; and (v) quantitative and qualitative behaviour (*cmp*. Figure 2 and Table 1). Additional “meta-properties” and provenance information further improve the comparison (e. g. information about file format or year of development).

**Figure 2.**
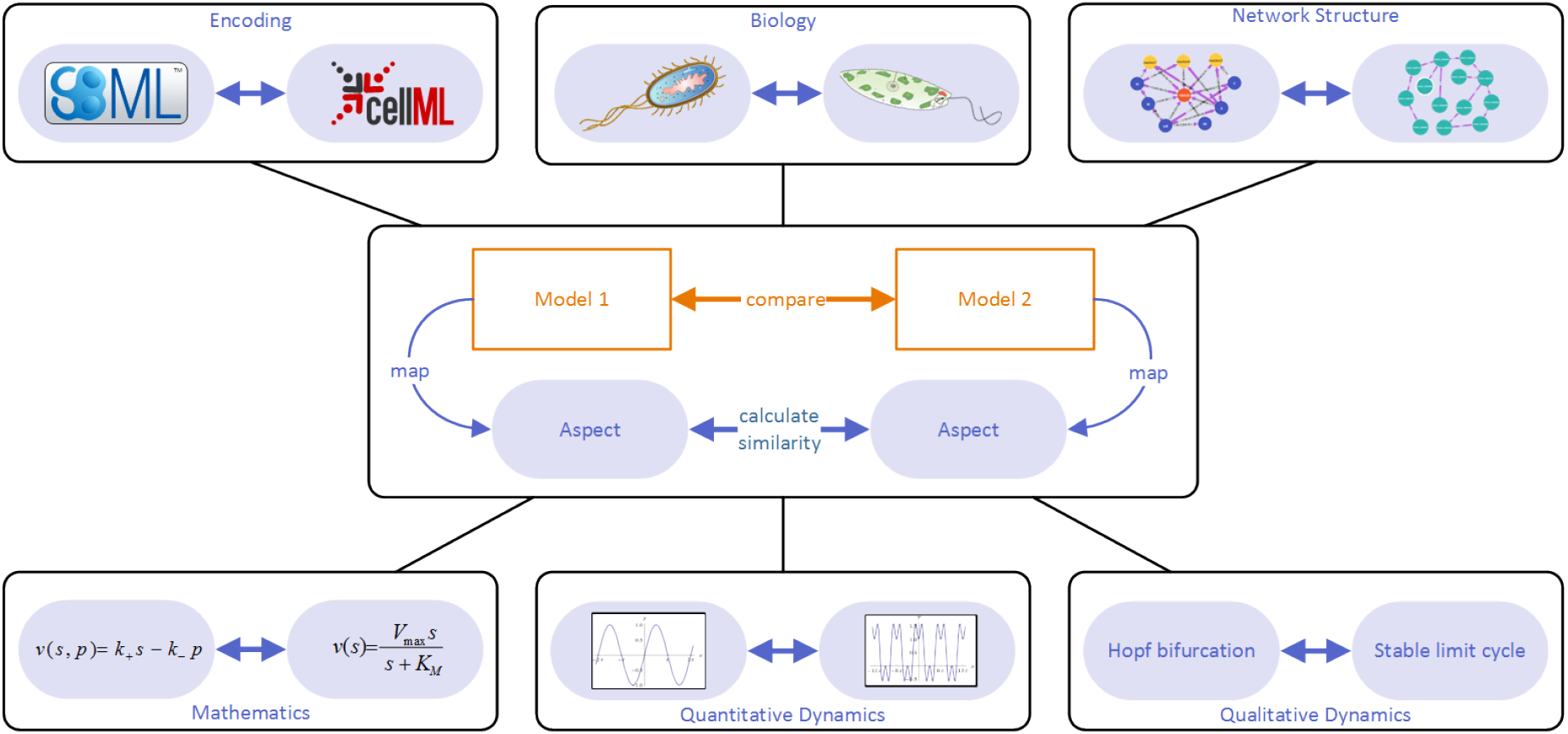
**Similarity measures for models can be derived from similarity measures for model aspects**. Automatic model comparison needs to rely on quantitative similarity measure. A practical way of defining such measures is to reduce both models to some relevant aspect, e. g., network structure, for which a similarity measure has already been defined. In an abstract scheme for model comparison, two models are mapped onto a specific focal aspect.

**Table 1.** Model aspects and related similarity measures. The table lists model aspects that are relevant for similarity, examples of similarity measures based on these aspects, examples of software tools supporting these types of similarity calculations, and practical application cases.

| Model aspect | Existing measures | Tool support (selection) | Applications |
| --- | --- | --- | --- |
| Encoding | Similarity on the XML-encoding (SBML or CellML), Levenshtein distance | BiVeS [19] Unix Diff | Model version control |
| Biological meaning | Similarity regarding biological meaning of model components | SemanticSBML, Semantic Measures Library and Toolkit Model set comparison, MORRE [20] | Search, retrieval, comparison of model intentions and assumptions |
| Network structure | Graph similarity, stoichiometric similarity | Cytoscape [21], graph libraries (e.g., JUNG <sup>1</sup>) | Model merging, extraction and comparison of submodels |
| Mathematical statements, quantitative and qualitative behaviour | Difference in statements and numbers; correlation between states, time-series, steady-states | Semantic Measures Library and Toolkit (for SBO annotations) | Compare dynamic behaviour; classification of models (e.g. oscillatory vs non-oscillatory glycolysis models) |

### 3.1 Model encoding

Models can be directly compared in their encoded form, i. e., in a specific file format. If models are encoded as computer programs, e. g. in MATLAB, a comparison on the syntactic level can be performed using tools for difference detection in software code (e.g., `diff`). Due to the lack of a predetermined structure of general computer programs, such a comparison does rarely yield satisfactory results. It is most useful for comparing versions of the same model. However, standardised formats such as SBML and CellML have a well-defined XML-syntax that restricts the number of possible operations. On the one hand, this constitutes a limitation in what can be encoded. On the other hand, the finite number of supported structures facilitates the implementation of domain-specific algorithms. These algorithms utilise the rich information about the encoded mechanisms, the components of models and their interactions, and the biological meaning of (parts of) the model. Similarly, simulations of models, and the context of the simulation, can be encoded in standard formats such as the Simulation Experiment Description Markup Language (SED-ML [22]). Thus the comparison of the models can convey information about their biological meaning and about their behaviour.

Several algorithms use this reduced approach of determining the similarity of models (or model versions) by comparing their syntactic structure, e. g. [19, 23, 24]. Some software tools compare representations of a model while taking the specific syntactic structure and its corresponding biological meaning into account, while others compare the representations directly and subsequently interpret the results with respect to the biology. For example, XML patches [23] are generated to compare models at the XML level only, while the BiVeS tool described in [19] also considers the structure of the actual model representation format.

### 3.2 Biological meaning

When aiming at a comparison of models with respect to the biology, the meaning of the components must be considered. The biological entities in a model (substances, reactions, cell compartments, etc.) must therefore be explicitly associated with a biological description. To this end, biological entities are usually semantically annotated with terms from biological and biomedical ontologies [25]. Model annotations may also refer to entries in biological databases from which ontology-based annotations can then be derived, e. g., UniProt [26].

Most models in open repositories are annotated with terms from ontologies such as the Gene Ontology (GO) [27], ChEBI [8], or the Systems Biology Ontology (SBO) [25]. Different ontologies cover different domains of knowledge and allow for model comparison related to these domains: model annotations referring to GO terms allow for defining similarity related to the biological function or cellular location of biological entities; annotations based on ChEBI allow for similarity based on chemical compounds; annotations based on the SBO allow for similarity based on biochemical rate laws or on the roles of chemical compounds in reaction mechanisms, etc. Similarity measures for models with respect to their semantic annotations can be defined on the basis of existing similarity measures for ontology elements [28]. These measures help to identify similarities in model matching and retrieval tasks [12, 29].

### 3.3 Network structure

Biological network structures, and especially their statistical properties, received large attention in systems biology research [30]. Network structures can be extracted from encoded models, and network alignments between models allow for detecting specific structural differences and similarities. Gay *et al*., for example, proposed graph-matching techniques to assess similarities between models [31]. They reduced the models to directed, bipartite reaction graphs and searched for epimorphisms between the graph structures. Subsequently, they evaluated the method on SBML models from BioModels. Graphs have also been used to measure the similarity of models based on the motifs they contain [32]. Motifs are small partial subgraphs that are statistically overrepresented in a network. Motifs are often interpreted as small functional units, and it has been shown that networks realizing similar tasks (e.g., signaling pathways) contain similar motifs [33]. Therefore, comparing the motif distributions in biological networks could provide information about typical functions that these networks perform.

### 3.4 Mathematical statements and model behaviour

Models can be compared by their quantitative or qualitative dynamic behaviour. This behaviour can either be determined from the mathematical structure or be observed in simulations. Similarity of behaviour has been associated with similar internal mechanisms and with dependencies on common extrinsic factors [16]. While we are not aware of any system that formally compares models with respect to dynamic behaviour, there are projects that compare, for a given model, the simulation results obtained from different simulators [34]. Other projects, such as the Functional Curation framework, compare the dynamic effects of particular simulated perturbations [35, 36]. A comparison of qualitative model behaviour (e.g. oscillation, steady state) can be useful to identify similarities between biological mechanisms. [16].

Annotations of reactions to terms in SBO may in future help to determine behavioural similarities. SBO is, for example, already used to annotate mathematic rate laws and therefore a good candidate for a qualitative comparison of mathematical expressions. If mathematical expressions are similar, then the models may also show a similar dynamic behaviour.

## 4 Practical applications of similarity measures

To profit from the wealth of published models, modellers need software that helps them explore, access, compare, simulate, and combine models with minimal effort. We distinguish two possible lines of action: On the one hand, researchers may either analyse small, defined sets of models, and classify or align them; as an example, Figure 3 shows how a dynamical cell cycle model could be compared to a simplified variant of the model from the same publication based on model files available at BioModels.net. On the other hand, researchers may search for models in databases such as BioModels, possibly starting from some query model of interest. Methods for model comparison are equally important in both cases. In the following, we describe basic use cases of model comparison as well as existing tools and methods devoted to these tasks.

**Figure 3.**
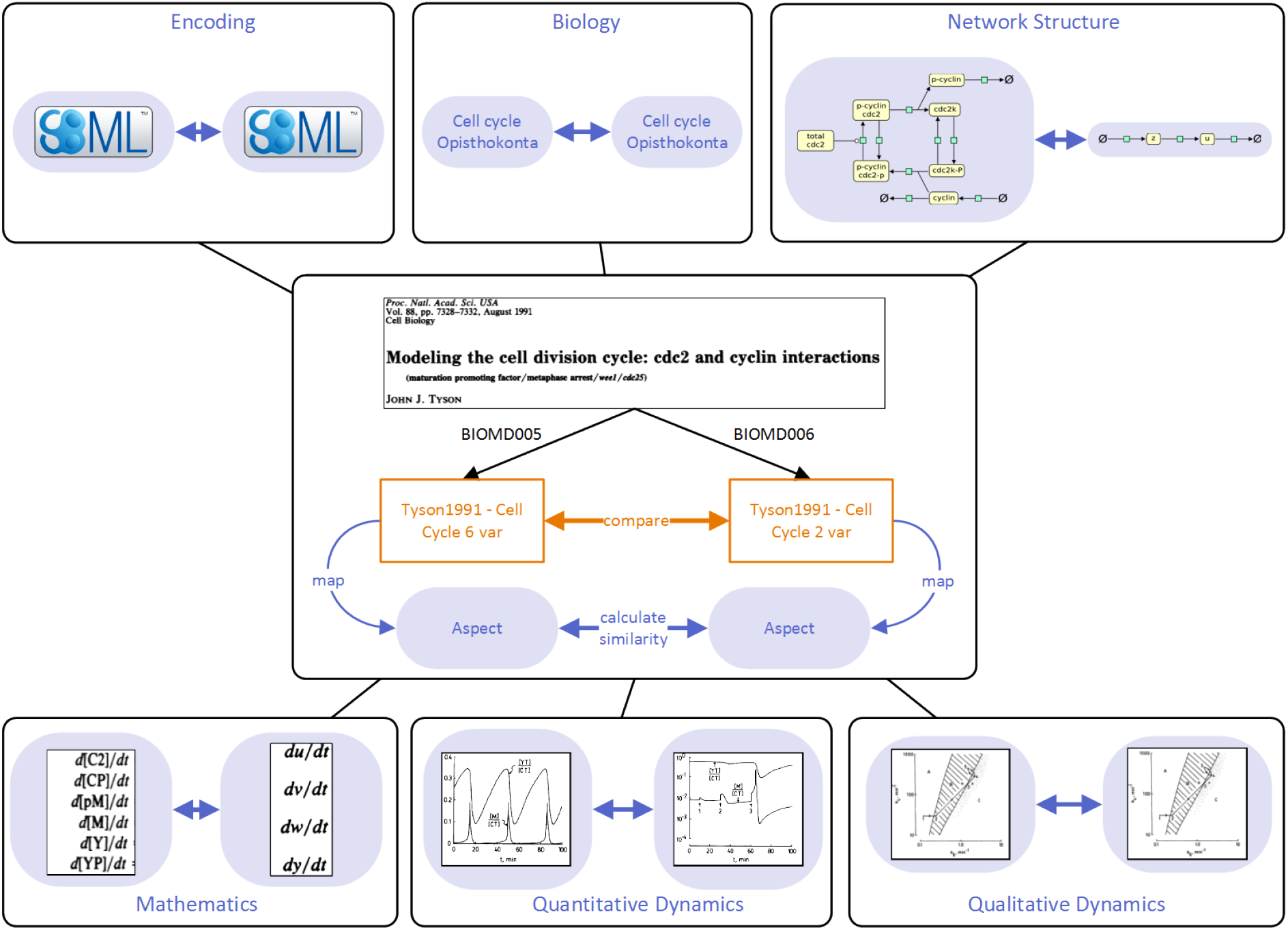
**Comparison of two models describing the cell division cycle**. We compare two dynamical models that stem from the same publication [37] and concern the action of a cyclin and a cyclin-dependent kinase. While the first model describes the phosphorylation of the two compounds by explicit reactions, the second model is simplified and captures only the general dynamics. The model files, obtained from BioModels (models BIOMD0000000005 and BIOMD0000000006), are encoded in SBML and carry the same semantic annotations, referring to the organism and biological pathway described. Due to their different levels of resolution, the models differ in their network structures and mathematical statements. The second model contains fewer variables and equations, which leads to a different quantitative dynamics. Nevertheless, the qualitative dynamic behaviour is the same given a suitable parametrisation. To automate all these comparisons, which would intuitively come to a modeller’s mind, the model aspects described above must be captured by similarity scores.

### 4.1 Model search and clustering

Number and sizes of available models are increasing beyond what even the most well-read scholar can review or analyse, as examplified in the growth of BioModels^2^. Probably the most common use of similarity measures is for search. Researchers query repositories to obtain models related to a given keyword, to a query model, or to a data set.

In a keyword search, a user enters a set of terms to retrieve models that match these terms. In the simplest form, a model can be represented by a “bag of terms”, i.e., a list of relevant keywords or annotations to which the user’s query terms can be matched. Similarity between query terms and model can then be defined using information retrieval measures. For example, a similarity score can be calculated from the frequency with which a term occurs, or from semantic similarity measures between terms [20]. Instead of such “bag-of-term” representations, models may also be represented in a structured form to incorporate network information, semantic annotations, and other associated meta-data. Keyword search is state-of-the-art in open model repositories. BioModels, for example, incorporates the aspects “model encoding” and “biological meaning” (*cf*. Table 1). A query by model encoding matches exactly a set of terms associated with a model. A query by biological meaning enables more sophisticated semantic searches. Search engines can be coupled with ranking algorithms to ensure that the most relevant search results appear in the top of the result list. The Physiome Model Repository (PMR2 [11]), for example, uses a Lucene-based ranking algorithm that incorporates model encoding, biological meaning, and other meta-data [12].

If the starting point of a search is a model, then the aim of a search is to find similar models. SemanticSBML [14] allows users to provide their own model as an input query. The search engine then finds SBML models that resemble the input model with respect to semantic annotations [29]. The search operates on the openly available models from BioModels.

Similarity measures can also be used to calculate the similarity between models and query terms. Based on functions for model ranking [12, 29], similarity measures can also help to cluster models into similar sets. For example, a cluster may organise models into thematic sets such as “models describing metabolism”, “the cell cycle”, or “models showing calcium oscillations”. Thematic model sets may be characterised through semantic annotations [38] or through recurring structural patterns [39]. BioModels, for example, implemented a web-based model browser that clusters models based on GO terms. In future, models may also be clustered based on biological motifs in their networks [40]. Thematic sets can be searched and compared more easily, e. g., when constructing comprehensive models.

### 4.2 Network alignments

Network alignments help to detect structural overlaps between pathways or networks. They are a basic tool in model merging tasks, and they can be used to define similarity scores. For example, an alignment of kinetic models of the metabolism showed that the mechanism is generally well covered. Parts of the central metabolism are even heavily overrepresented [29]. If coupled with semantics-based measures, network alignments can be used to score the relative overlap of networks. However, aligning two networks may be challenging if only few of the components are precisely annotated. One approach to address this problem is through semantic propagation [41], a method that combines semantic information with information about the model’s structure (e.g., reaction network, cellular compartments) to infer missing annotations and align networks at the same time. Interestingly, the method not only compares model components by their own annotations, but also by annotations of related components. For example, reactions can be compared by annotations of their reactants, and cell compartments can be compared by the compounds they contain.

### 4.3 Model version control

New insights about a biological system may call for an adjustment of network structure, mathematical formulae, or parameters of a model. Strong adjustments usually result in new model versions. Another frequent reason for updates are error corrections. The comparison of model versions can help to keep track of the model’s evolution in time, and it identifies points at which a model underwent major changes [42]. BiVeS [19] is a software library that aligns the XML encodings of two model versions, identifies and interprets changes, and measures the changes’ impact. BiVeS also considers characteristics of the model encoding format and differentiates between SBML and CellML. Single diffs are automatically annotated to terms of COMODI, an ontology describing possible changes on computational biology models [43].

## 5 Discussion

Many efforts were made in the past years to improve the reusability of computational biology models as well as the reproducibility of associated results [44, 45, 46]. Model repositories collect curated models ready to be reused, provide semantic annotations, and offer instructions on how to simulate the models appropriately. With standardised, annotated models being available, automatic model comparison has become a feasible task. One important application is the search for models. Curated models in standard formats can also be version controlled, allowing researchers to follow a model’s evolution through time. Key challenges in model search and version control include an appropriate ranking of search results, a measure for the quality of the retrieved models, and a framework to implement full model provenance.

Similarity measures are valuable beyond standard model management tasks. They are crucial for merging existing pathway models into larger cell models. The map of Human Metabolism developed in the ReconX project, for instance, relies strongly on previously published models [47]. When merging two models, parts of their networks may have to be exchanged or replaced by suitable alternatives, requiring tools that can compare both, models and model parts, and find the most similar matches. Since these tools need to solve similar problems and share similar difficulties, we propose to study model similarity as a general task. As a key challenge, we identify the appropriate choice of the similarity measure: How can we define computational measures that reflect user’s expectations about similarity? And how can these measures, with various criteria, be applied in complex tasks such as search or merging? A number of issues need to be addressed in future research to create a general framework for model similarity and model comparison.

### Implement similarity measures for all model aspects

When comparing models, current software focuses on two of the aspects defined in this paper, namely the biological meaning and the model encoding. Other model aspects are not yet commonly used. However, their implementation in existing algorithms is feasible and will improve model comparison. Dynamic behaviour, obtained from model simulations, could reveal similarities between biological processes during execution. These similarities will not show when considering pathway structure alone. One can directly compare simulated time series (as showcased by the Cardiac Physiology Web Lab [36]), or build a system that compares their semantic annotations with quantitative or qualitative behaviour observed in simulations. A valuable resource of terms for dynamic behaviour is the TEDDY ontology (TErminology for the Descriptions of DYnamics, [25]). Likewise, improved similarity measures could be obtained by more extensively exploring graph matching and graph similarity algorithms [48, 49], as suggested in [50]. Comparison of network structure can help with matching dynamical models to experimentally determined interaction networks. Equally, mathematical expressions could be compared directly to yield deeper insights into similarities of the models’ behaviour. Furthermore, information that is related to the model may become relevant, such as the purpose of an investigation or the modellers’ intentions. However, these information must first be formalised – a new and interesting challenge.

### Combine similarity measures

Today’s software mostly compares models on a single model aspect. Since different similarity measures have proven useful for specific applications, we expect that the combination of aspects would enable an even more powerful comparison of models. Consider the following example: A scientist searches for a MAP kinase cascade model that contains regulatory feedback loops and shows dynamic oscillations. In a first step, the search tool could search for models on the specific biological system (using semantic comparison). Then, it could filter the intermediate results for specific network topologies. Finally, a second filter could be applied to select models with oscillatory behaviour. If documentation in a repository is complete, a behaviour-based comparison of the remaining models could be performed (e.g., by evaluating associated simulation descriptions in SED-ML format, and by comparing TEDDY terms therein). Eventually, an overall similarity joining the different aspects could be defined. However, such a procedure requires suitable functions for combining the individual similarities in a single formula. Furthermore, it should be possible for users to specify weights, i. e. a relative importance, for each of the aspects.

### Improve software support

To allow for extended similarity measures and to integrate them easily into software applications, new tools need to be developed. Interoperability through standard formats and common libraries must be ensured. Such tools could incorporate a large set of existing similarity measures and provide functions for projecting models onto their relevant aspects. In addition, the scientific community has developed shared resources for models and associated metadata, widely accessible through graph databases [20] or publicly accessible Semantic Web resources [51]. Tools to process models and determine similarity should be able to access and interoperate with these shared resources.

### Provide tools to align models and data by similarity

We discussed how similarity measures link models to other models or queries. Another exiting approach is to link models to experimental data sets. We envision that a comparison between models and other datasets is possible through projection onto a common aspect. For example, both models and a patient’s cancer genomics dataset can be projected to common proteins. Subsequently, semantic annotations of biological processes can be compared between the model entities and the data items. Afterwards, similarity measures for biological meaning can help to identify whether observations in the patient match a particular state predicted by a model.

### Develop intuitive user interfaces

When offering purpose-driven similarity measures, it is important to communicate the details of the scores to the users. Consequently, there is a need to develop clear and intuitive user interfaces that show both, the results of a similarity score and the details of calculation. For example, a system that supports ranked retrieval should return a ranked list of models, with a detailed description of filtering processes and relationships between models and query. Some software tools already visualise element alignments between models as network graphs (e. g., SemanticSBML, STON, BudHat [42]), or present ranking scores for retrieved models (e.g., SemanticSBML [29], MASYMOS [20]). However, an explanation of the steps leading to the similarity scores will increase the trust of the users and support decisions for a particular search result.

### 5.1 Conclusions

Similarity between models is assessed by a variety of software applications. Here, we classified and reviewed model similarity measures systematically. A number of aspects help with determining the similarity between models: the model encoding, when comparing versions of a model; the mathematical description of a model, when investigating the systems’ dynamic behaviour; the biological elements appearing in a model, when searching for models of a specific biological system or phenomenon; the network structure, when investigating the reuse of models as submodels in large networks; the parameter values in a model, during functional curation; and simulation outcomes, when comparing behaviour and sensitivity of a model. We envision a general framework for model similarity, based on a systematic treatment of models and their aspects. Such a framework will enhance the automated processing of models and will have numerous applications in computational systems biology.

## 6 Acknowledgments

The authors would like to thank the participants of the 2013 Model Meeting in Rostock (Germany) for valuable discussions on similarity notions of models. The meeting was funded through the BMBF e:Bio program, grant no. FKZ0316194. DW is funded through the Junior Research Group SEMS, BMBF e:Bio program, grant no. FKZ0316194. TK is funded through the BMBF via the Greifswald Approach to Individualized Medicine (GANI_MED) (grant 03IS2061A) and “Unternehmen Region” as part of the ZIK-FunGene (grant 03Z1CN22).

## Footnotes

^1^http://jung.sourceforge.net/index.html

^2^BioModels stores 144,366 models as of April 16^th^ 2015

## References

[1] N. Le Novère. Quantitative and logic modelling of molecular and gene networks. Nature Reviews Genetics, 2015.

[2] O. Wolkenhauer. Why model? Frontiers in physiology, 5, 2014.

[3] R. Rosen. Life itself. Columbia University Press, New York, 1991.

[4] R.-S. Wang, A. Saadatpour, and R. Albert. Boolean modeling in systems biology: an overview of methodology and applications. Physical biology, 9(5):055001, 2012.

[5] M. Hucka, A. Finney, H.M. Sauro, H. Bolouri, J.C. Doyle, H. Kitano, A.P. Arkin, B.J Bornstein, D. Bray, A. Cornish-Bowden, A.A. Cuellar, S. Dronov, E.D. Gilles, M. Ginkel, V. Gor, I.I. Goryanin, W.J Hedley, T.C. Hodgman, J.S Hofmeyr, P.J. Hunter, N.S Juty, J.L. Kasberger, A. Kremling, U. Kummer, N. Le Novère, L.M. Loew, D. Lucio, P. Mendes, E. Minch, E.D. Mjolsness, Y. Nakayama, M.R. Nelson, P.F Nielsen, T. Sakurada, J.C. Schaff, B.E Shapiro, T.S. Shimizu, H.D Spence, J. Stelling, K. Takahashi, M. Tomita, J.M. Wagner, and J. Wang. The systems biology markup language (SBML): a medium for representation and exchange of biochemical network models. Bioinformatics, 19(4):524–531, 3 2003.

[6] A.A. Cuellar, C.M. Lloyd, P.F. Nielsen, D.P. Bullivant, D.P. Nickerson, and P.J. Hunter. An Overview of CellML 1.1, a Biological Model Description Language. SIMULATION, 79(12):740–747, 2003.

[7] M. Hucka, F.T. Bergmann, S.M Keating, and L.P. Smith. A profile of today's SBML-compatible software. In e-Science Workshops (eScienceW), 2011 IEEE Seventh International Conference on, pages 143–150. IEEE, 2011.

[8] K. Degtyarenko, P. de Matos, M. Ennis, J. Hastings, M. Zbinden, A. McNaught, R. Alcántara, M. Darsow, M. Guedj, and M. Ashburner. ChEBI: a database and ontology for chemical entities of biological interest. Nucleic Acids Research, 36(suppl 1):D344–D350, 2008.

[9] S.A. Sansone, P. Rocca-Serra, D. Field, F. Maguire, C. Taylor, O. Hofmann, et al. Toward interoperable bioscience data. Nature Genetics, 44(2):121–126, 2012.

[10] D. Waltemath. Management of simulation studies in computational biology. In Invited presentations, junior research groups and research highlights at GCB 2015. PeerJ preprints, 2015.

[11] T. Yu, C.M. Lloyd, D.P. Nickerson, M.T. Cooling, A.K. Miller, A. Garny, J.R. Terkildsen, J. Lawson James, R.D. Britten, P.J Hunter, et al. The physiome model repository 2. Bioinformatics, 27(5):743–744, 2011.

[12] R. Henkel, L. Endler, A. Peters, N. Le Novère, and D. Waltemath. Ranked retrieval of computational biology models. BMC Bioinformatics, 11(1):423, 2010.

[13] M. Lange, R. Henkel, W. Müller, D. Waltemath, and S. Weise. Information retrieval in life sciences: a programmatic survey. In Approaches in Integrative Bioinformatics, pages 73–109. Springer, 2014.

[14] F. Krause, J. Uhlendorf, T. Lubitz, M. Schulz, E. Klipp, and W. Liebermeister. Annotation and merging of SBML models with semanticSBML. Bioinformatics, 26(3):421–422, 2010.

[15] C. Clark and J. Kalita. A comparison of algorithms for the pairwise alignment of biological networks. Bioinformatics, 30(16):2351–2359, 2014.

[16] A. Tapinos and P. Mendes. A method for comparing multivariate time series with different dimensions. PloS one, 8(2):e54201, 2013.

[17] R.O. Duda, P.E. Hart, and D.G. Stork. Pattern Classification. Wiley, 2nd, 2000.

[18] N. Le Novère, M. Hucka, H. Mi, S. Moodie, F. Schreiber, A. Sorokin, E. Demir, K. Wegner, M.I. Aladjem, S.M Wimalaratne, F.T. Bergman, R. Gauges, P. Ghazal, H. Kawaji, L. Li, Y. Matsuoka, A. Villéger, S.E. Boyd, L. Calzone, M. Courtot, U. Dogrusoz, T.C. Freeman, A. Funahashi, S. Ghosh, A. Jouraku, S. Kim, F. Kolpakov, A. Luna, S. Sahle, E. Schmidt, S. Watterson, G. Wu, I. Goryanin, D.B. Kell, C. Sander, H. Sauro, J.L. Snoep, K. Kohn, and H. Kitano. The systems biology graphical notation. Nature Biotechnology, 27(8):735–741, 8 2009.

[19] Martin Scharm, Olaf Wolkenhauer, and Dagmar Waltemath. An algorithm to detect and communicate the differences in computational models describing biological systems. Bioinformatics, btv484, 2015.

[20] R. Henkel, O. Wolkenhauer, and D. Waltemath. Combining computational models, semantic annotations and simulation experiments in a graph database. Database, 2015:bau130, 2015.

[21] Paul Shannon, Andrew Markiel, Owen Ozier, Nitin S Baliga, Jonathan T Wang, Daniel Ramage, Nada Amin, Benno Schwikowski, and Trey Ideker. Cytoscape: a software environment for integrated models of biomolecular interaction networks. Genome research, 13(11):2498–2504, 2003.

[22] D. Waltemath, R. Adams, F.T. Bergmann, M. Hucka, F. Kolpakov, A.K. Miller, I.I. Moraru, D. Nickerson, S. Sahle, J.L. Snoep, et al. Reproducible computational biology experiments with SED-ML - the simulation experiment description markup language. BMC systems biology, 5(1):198, 2011.

[23] P. Saffrey and R. Orton. Version control of pathway models using XML patches. BMC Systems Biology, 3(1):34, 2009.

[24] A. Miller, T. Yu, R. Britten, M. Cooling, J. Lawson, D. Cowan, A. Garny, M. Halstead, P. Hunter, D. Nickerson, G. Nunns, S. Wimalaratne, and P.F. Nielsen. Revision history aware repositories of computational models of biological systems. BMC Bioinformatics, 12(1):22, 2011.

[25] M. Courtot, N. Juty, C. Knüpfer, D. Waltemath, A. Zhukova, A. Dräger, M. Dumontier, A. FinneyA, M. Golebiewski, J. Hastings, S. Hoops, S. Keating, D.B. Kell, S. Kerrien, J. Lawson, A. Lister, J. Lu, R. Machné, P. Mendes, M. Pocock, N. Rodriguez, A. Villeger, D.J. Wilkinson, S. Wimalaratne, C. Laibe, M. Hucka, and N Le Novère. Controlled vocabularies and semantics in systems biology. Molecular Systems Biology, 7, 2011.

[26] R. Apweiler, A. Bairoch, C.H. Wu, W.C Barker, B. Boeckmann, S. Ferro, E. Gasteiger, H. Huang, R. Lopez, M. Magrane, M.J. Martin, D.A Natale, C. O'Donovan, N. Redaschi, and L.S. Yeh. UniProt: the universal protein knowledgebase. Nucleic Acids Research, 32(D1):115–119, 1 2004.

[27] M. Ashburner, C.A. Ball, J.A Blake, D. Botstein, H. Butler, J.M. Cherry, A.P Davis, K. Dolinski, S.S. Dwight, J.T Eppig, M.A. Harris, D.P Hill, L. Issel-Tarver, A. Kasarskis, S. Lewis, J.C. Matese, J.E Richardson, M. Ringwald, G.M. Rubin, and G. Sherlock. Gene Ontology: tool for the unification of biology. Nature Genetics, 25:25–29, 2000. The Gene Ontology Consortium.

[28] C. Pesquita, D. Faria, A.O. Falcao, P. Lord, and F.M. Couto. Semantic similarity in biomedical ontologies. PLoS Computational Biology, 5(7):e1000443, 07 2009.

[29] M. Schulz, F. Krause, N. Le Novère, E. Klipp, and W. Liebermeister. Retrieval, alignment, and clustering of computational models based on semantic annotations. Molecular systems biology, 7(1), 2011.

[30] U. Alon. An introduction to systems biology: design principles of biological circuits. CRC press, 2006.

[31] S. Gay, S. Soliman, and F. Fages. A graphical method for reducing and relating models in systems biology. Bioinformatics, 26(18):i575–i581, 2010.

[32] R. Milo, S. Shen-Orr, S. Itzkovitz, N. Kashtan, D. Chklovskii, and U. Alon. Network motifs: simple building blocks of complex networks. Science, 298(5594):824–827, 2002.

[33] U. Alon. Biological networks: the tinkerer as an engineer. Science, 301(5641):1866–1867, 2003.

[34] F.T. Bergmann and H.M. Sauro. Comparing simulation results of SBML capable simulators. Bioinformatics, 24(17):1963–1965, 2008.

[35] J. Cooper, G.R. Mirams, and S.A. Niederer. High-throughput functional curation of cellular electrophysiology models. Progress in biophysics and molecular biology, 107(1):11–20, 2011.

[36] J. Cooper, M. Scharm, and G.R. Mirams. The cardiac electrophysiology web lab. Biophysical Journal, 110(2):292–300, 2016.

[37] J.J. Tyson. Modeling the cell division cycle: cdc2 and cyclin interactions. Proceedings of the National Academy of Sciences, 88(16):7328–7332, 1991.

[38] R. Alm, D. Waltemath, M. Wolfien, O. Wolkenhauer, and R. Henkel. Annotation-based feature extraction from sets of SBML models. Journal of Biomedical Semantics, 6(1):20, 2015.

[39] R. Henkel, F. Lambusch, and D. Waltemath. Finding pattern in biochemical reaction networks-a sub-graph mining approach. PeerJ PrePrints, 3:e1848.

[40] J.J. Tyson and B. Novák. Functional motifs in biochemical reaction networks. Annual review of physical chemistry, 61:219, 2010.

[41] M. Schulz, E. Klipp, and W. Liebermeister. Propagating semantic information in biochemical network models. BMC Bioinformatics, 13(1):18, 2012.

[42] D. Waltemath, R. Henkel, R. Hälke, M. Scharm, and O. Wolkenhauer. Improving the reuse of computational models through version control. Bioinformatics, 29(6):742–748, 2013.

[43] Martin Scharm, Dagmar Waltemath, Pedro Mendes, and Olaf Wolkenhauer. Comodi: An ontology to characterise differences in versions of computational models in biology. 4:e1857v1, 2016.

[44] F. Krause, M. Schulz, N. Swainston, and W. Liebermeister. Methods in Systems Biology, 500 Methods in Enzymology, chapter Sustainable model building: the role of standards and biological semantics, pages 371–395. 2011.

[45] D. Waltemath, R. Henkel, F. Winter, and O. Wolkenhauer. Reproducibility of model-based results in systems biology. In Systems Biology, pages 301–320. Springer, 2013.

[46] D. Waltemath and O. Wolkenhauer. How modeling standards, software, and initiatives support reproducibility in systems biology and systems medicine. under revision.

[47] I. Thiele, N. Swainston, R.M.T. Fleming, A. Hoppe, S. Sahoo, M.K. Aurich, H. Haraldsdottir, M.L. Mo, O. Rolfsson, M.D. Stobbe, et al. A community-driven global reconstruction of human metabolism. Nature Biotechnology, 31(5):419–425, 2013.

[48] J. Berg and M. Lässig. Local graph alignment and motif search in biological networks. Proceedings of the National Academy of Sciences of the United States of America, 101(41):14689–14694, 2004.

[49] X. Yan and J. Han. gspan: Graph-based substructure pattern mining. In Data Mining, 2002. ICDM 2003. Proceedings. 2002 IEEE International Conference on, pages 721–724. IEEE, 2002.

[50] C. Rosenke and D. Waltemath. How can semantic annotations support the identification of network similarities? In SWAT4LS, 2014.

[51] S. Jupp, J. Malone, J. Bolleman, M. Brandizi, M. Davies, L. Garcia, A. Gaulton, S. Gehant, C. Laibe, N. Redaschi, et al. The ebi rdf platform: linked open data for the life sciences. Bioinformatics, 30(9):1338–1339, 2014.

